# The impact of coregistration of gradient recalled echo images on quantitative susceptibility and R2* mapping at 7T

**DOI:** 10.1101/529891

**Authors:** Seongjin Choi, Xu Li, Daniel M. Harrison

**Author notes:** Corresponding author: Seongjin Choi, (SC). **Grant support:** This study was funded through grants from Bayer Neurosciences, NIH/NIBIB P41EB015909 and NINDS 1K23NS072366.

## Abstract

**Introduction:** While image coregistration is inevitable and quantitative parametric maps, such as R2* and QSM, are increasingly used in multi-parametric studies of the brain, there is a lack of investigations on the reliability of quantitative metrics after coregistration. The purposes of this study were 1) to evaluate the reliability of R2* and QSM at 7T and 2) to assess the statistical agreement in the quantitative metrics obtained by two different coregistration approaches.

**Methods:** We compared the reliability of R2* and quantitative susceptibility maps obtained from brains of eight healthy participants by two coregistration approaches: 1) transformation of pre-processed quantitative maps and 2) processing quantitative maps after transformation using pixel- and ROI-based analyses. Two-sample Kolmogorov-Smirnov, Mann-Whitney U, Paired T, Intraclass-correlation tests were performed appropriately.

**Results:** R2* remained invariant regardless of the cogeneration timing. However, magnetic susceptibility was significantly altered when processed in transformed space, whereas it remained invariant in all KS-tests and pixel values were only different in 2 out of 64 U-tests for direct QSM coregistration. Paired t-test revealed that ROI-based group-mean R2* was invariant to both approaches, while group-mean susceptibility was invariant to direct coregistration but differed in one structure processed in transformed space. For all pairs of measurements of R2*, ICCs were excellent. ICCs for magnetic susceptibility were excellent when processed in its native space while the ICCs were lower than 0.9 or poor when processed in transformed space. Further analysis revealed that the choice of interpolation approach affected the resultant QSM.

**Conclusions:** Our study shows that R2* could be safely processed in a transformed space, whereas QSM was less reliable when processed in the transformed space. Hence, caution is advised when using QSM in a multi-parametric study, and it is strongly recommended to process QSM in its native space prior to any coregistration or spatial transformation.

## Introduction

Image coregistration is an inevitable step of data processing in contemporary studies using magnetic resonance imaging (MRI) data because of increasing demands for multi-parametric image analyses. Further, combined quantitative image analysis often requires coregistration of several MR images of various spatial resolutions as well as different contrasts into a common image space. Image coregistration is also essential in longitudinal studies, in which MR images are acquired and accumulated across a few time points.

Image coregistration often employs multiple sophisticated mathematical algorithms, for example, geometric transformation, interpolation, and estimation of pixel values (PVs) from a source image in a transformed space. Hence, in the transformed space, the resulting image may consist of voxels of altered spatial locations, dimensions, and intensity values compared to the original image. Therefore, some researchers may raise concerns about the reliability of metrics derived from the altered PVs in the transformed space.

Although individual PVs of the coregistered image in a transformed space may be altered, data derived from a post-registration image may still be useful for multi-parametric analyses if the descriptive statistics and the distribution profile of the PVs in structural labels or regions of interest (ROI) remain statistically identical. If the statistical features of altered PVs in the transformed space are significantly different from those of its source MR image, any metrics derived from the transformed image are not reliable and should not be considered in further analysis.

While image coregistration is inevitable and various quantitative parametric maps, such as R2* and quantitative susceptibility mapping (QSM), are increasingly used in studies employing multiparametric assessment of the brain, there is a lack of investigations on the reliability of the derived quantitative metrics after source image coregistration. There are a limited number of studies investigating the reproducibility of QSM. [1–4] Hinoda et al. assessed the consistency and reproducibility of QSM at 3T and 1.5T and demonstrated good reproducibility of QSM processed by two different background phase removal algorithms at both field strengths. [1] Deh et al. assessed the reproducibility of nonlinear MEDI-based QSM measurements in both control and patient groups for scanners from two different vendors and at both 3T and 1.5T. Their QSM measurements showed good intra-scanner and inter-scanner reproducibility for healthy and multiple sclerosis participants. [2] Santin et al. investigated the reproducibility of QSM measurements at 3T in healthy volunteers within and between different MR sessions using four different combinations of QSM reconstruction methods. [3] They also investigated the reproducibility of R2* measurements in comparison with QSM measurements. Lauzon et al. compared QSM acquired by various methodologies. [4] They showed that readout gradient polarity and accelerated parallel imaging did not alter the estimated susceptibility at 3T. In summary, these studies reported inter-vendor, inter-acquisition methods, inter/intra-scanner, inter-field (1.5 and 3T), and inter-QSM reconstruction methods agreements. More recently, a few more studies explored QSM reliability or reproducibility at ultra-high field (7T). Lancione et al. demonstrated excellent intra-scanner repeatability and inter-scanner reproducibility of QSM at both 3T and 7T within five healthy subjects of mixed gender. [5] Rua et al. investigated the reproducibility of QSM and R2* map at 7T and showed promising results for multi-center and multi-vendor planforms within three healthy male subjects. [6] Okada et al. investigated test-retest reliability of cortical R2* and QSM at 7T within sixteen healthy subject and both measures showed repeatable results in the wide cortical area except at the frontal-temporal base area. [7]

However, none of these studies investigated the impact of co-registration of source gradient recalled echo (GRE) images to a non-native space on the resulting parametric maps of R2* and quantitative susceptibility. In these studies, when inter-space coregistrations were required, the resultant quantitative maps were coregistered after being processed in their native spaces, although there has been no empirical evidence to do so. Moreover, reliabilities of QSM and R2* maps have not been fully explored at ultra-high field (7T). In this study, we investigated the influence of two different coregistration approaches on quantitative measurements (R2* and magnetic susceptibility) acquired at 7T in particular. We examined the reliability of quantitative measures collected within four atlas-based structural labels in deep brain areas bilaterally using both pixel- and ROI-based analyses. We addressed how two coregistration approaches differently impact on R2* map and QSM. Furthermore, we addressed how an insignificant modification of phase-image resolution led to a significant change in resultant QSM.

## Materials and Methods

This study consists of two stages. Initial stage was performed to assess the reliability of 7T R2* and QSM and impact of the coregistration approaches. Then, based on the initial analysis results, a further analysis was conducted to address the questions raised in the initial analysis.

### Participants

Eight healthy subjects with ages ranging from 32 to 57 (four men, four women) participated in the study. The Institutional Review Boards at the Johns Hopkins University School of Medicine and the Kennedy Krieger Institute approved protocols, and all participants provided written informed consent.

### Acquisition

MRI was acquired using a whole-body 7T Achieva MR scanner (Philips Medical Systems, Cleveland, OH, USA) and a volume-transmit, 32-channel receive head coil (Nova Medical, Wilmington, MA, USA). MRI protocol consisted of a magnetization prepared rapid acquisition gradient echo sequence (MPRAGE, TR/TE = 4.1/1.83 ms, sagittal acquisition, FOV = 180 × 220 × 220 mm^3^, image matrix = 180 × 220 × 220, flip angle = 7 degrees, TFE factor = 352, SENSE factor = 2 × 1 × 2) and a multi-echo 3D gradient-recalled echo sequence (GRE, TR/TE1/ΔTE = 68/4/2 ms, 8 echoes, axial acquisition, FOV = 220 × 220 × 110 mm^3^, image matrix = 224 × 224 × 110) with both magnitude and phase output.

### Image processing

#### Preprocessing

All pre- and post-processing steps were performed on a Linux (Debian 8) desktop with Neurodebian packages [8]. Before starting any image processing, all MRI data saved in Par/Rec format (Philips native format) were converted to NIfTI format using the ‘parrec2nii’ module in ‘NIPY’ package written in Python script language [9]. MPRAGE data were corrected for the bias field using the N4 bias-field correction algorithm prior to linear registration [10]. Then, linear coregistration of GRE data to MPRAGE space was performed using FLIRT (FMRIB, Oxford University) with rigid body transformation [11, 12]. A transformation matrix (GRE-to-MPRAGE transformation matrix) was acquired for coregistration of the first-echo magnitude image of GRE to MPRAGE space. The first-echo magnitude was chosen because it has the least signal loss due to intra-voxel spin dephasing and ensures preserving the most brain tissues. For calculating QSM in MPRAGE space, magnitude and phase image pairs were then converted to real and imaginary pairs using FSLUTILS included in FSL. The transformation matrix was then applied to the real and imaginary image pairs. Finally, the transformed real and imaginary pair was converted back to a magnitude and phase image pair for further processing in MPRAGE space. The QSM processing in a transformed space requires consideration of its dependency on coregistration transformation matrix. Hence, we assured use of a transformed dipole kernel in QSM processing in MPRAGE space by reflecting angle changes introduced by transformation matrix from GRE to MRPAGE spaces.

#### R2* mapping

Two sets of R2* maps were calculated in a voxel-wise manner using the power method, in which the decay of squared magnitudes from all eight echoes was fitted to a monoexponential model. [13] This was performed using an in-house tool written in MatLab (Mathworks, Inc., Natick, MA). One set of R2* maps was calculated in its native (GRE) space, and another set was calculated from GRE data transformed to MPRAGE space. The third set of R2* maps was acquired by direct transformation of the R2* map in its native space to MPRAGE space using the GRE-to-MPRAGE transformation matrix acquired during preprocessing.

#### Quantitative Susceptibility Mapping (QSM)

Three sets of QSM data were also calculated for each participant. The first two sets of QSMs were processed using GRE images in native space and GRE images transformed to MPRAGE space. Although QSM can be calculated from both single-echo and multi-echo GRE data [14], we only processed multi-echo QSM. QSM processing was accomplished using an in-house tool (JHU QSM Toolbox) written in Matlab. The first three echoes (at TE of 4 ms, 6 ms, and 8 ms) were excluded from calculating QSM to avoid non-linear temporal phase evolution in white matter when TE is shorter than 10 ms at ultra-high field [15, 16], and the other five echoes (with TE of 10 ms to 18 ms) underwent the phase processing and were averaged to generate the final frequency map and susceptibility maps. Phase images underwent a sequential process of phase unwrapping, background-field removal, and dipole inversion. Phase unwrapping employed a Laplacian-based phase unwrapping algorithm [17]. The unwrapped phase images were then divided by 2**π**\*TE to obtain an image of frequency shift in Hz. Background field removal was peformed by the V-SHARP method [18–20] with a maximum spherical kernel size of 6 mm and TSVD threshold of 0.05. Then, dipole inversion was performed by applying a modified structural feature based collaborative reconstruction (SFCR) algorithm with only L1-norm based regularization i.e. *λ*_1_ = *γ*_1_ = 100 and dropped L2-norm regularization i.e. *λ*_2_ = *γ*_2_ = 0 [21]. Finally, the last set of QSM was acquired by directly coregistering QSM in native (GRE) space to MPRAGE space using the GRE-to-MPRAGE transformation matrix saved in preprocessing.

#### Structural label preparation and data collection

Two sets of structural labels in MPRAGE and GRE spaces for each participant were prepared. Atlas-based structural labels were used to avoid biases introduced by manual drawing. We used structural labels obtained from the Harvard-Oxford atlas, which is included in FSL and based on T1-weighted images of 37 healthy subjects of mixed genders in standard MNI 152 space. The source atlas contained labels for each structure bilaterally. Four anatomical structures from this atlas were bilaterally selected for analysis: thalamus (TH_L and TH_R), caudate nucleus (CN_L and CN_R), globus pallidus (GP_L and GP_R), and putamen (PUT_L and PUT_R). Using FSL, the atlas-based labels were eroded by one voxel (with 3-D kernel) to ensure that measurements were performed only within the intended structures after the following image transformations and to reduce potential partial volume effect between other structures and other tissue types in the vicinity. The eroded structural labels were then inversely warped to the MPRAGE space of each participant using FNIRT available in FSL (FMRIB, Oxford University) [22]. These labels were used to collect PVs from parametric maps in MPRAGE space. Finally, the labels were linearly transformed to each participant’s GRE space with the inverse of the GRE-to-MPRAGE transformation matrix. These labels were used to collect PVs from parametric maps in GRE space. Using the two sets of labels in GRE and MPRAGE spaces, we sampled voxel-based data for the selected four structures bilaterally. These pixel-based data were further used to calculate group-mean values for each structural label. Fig 1 shows the eight structures in the standard (MNI), GRE, and MPRAGE spaces of a representative participant.

**Fig 1.**
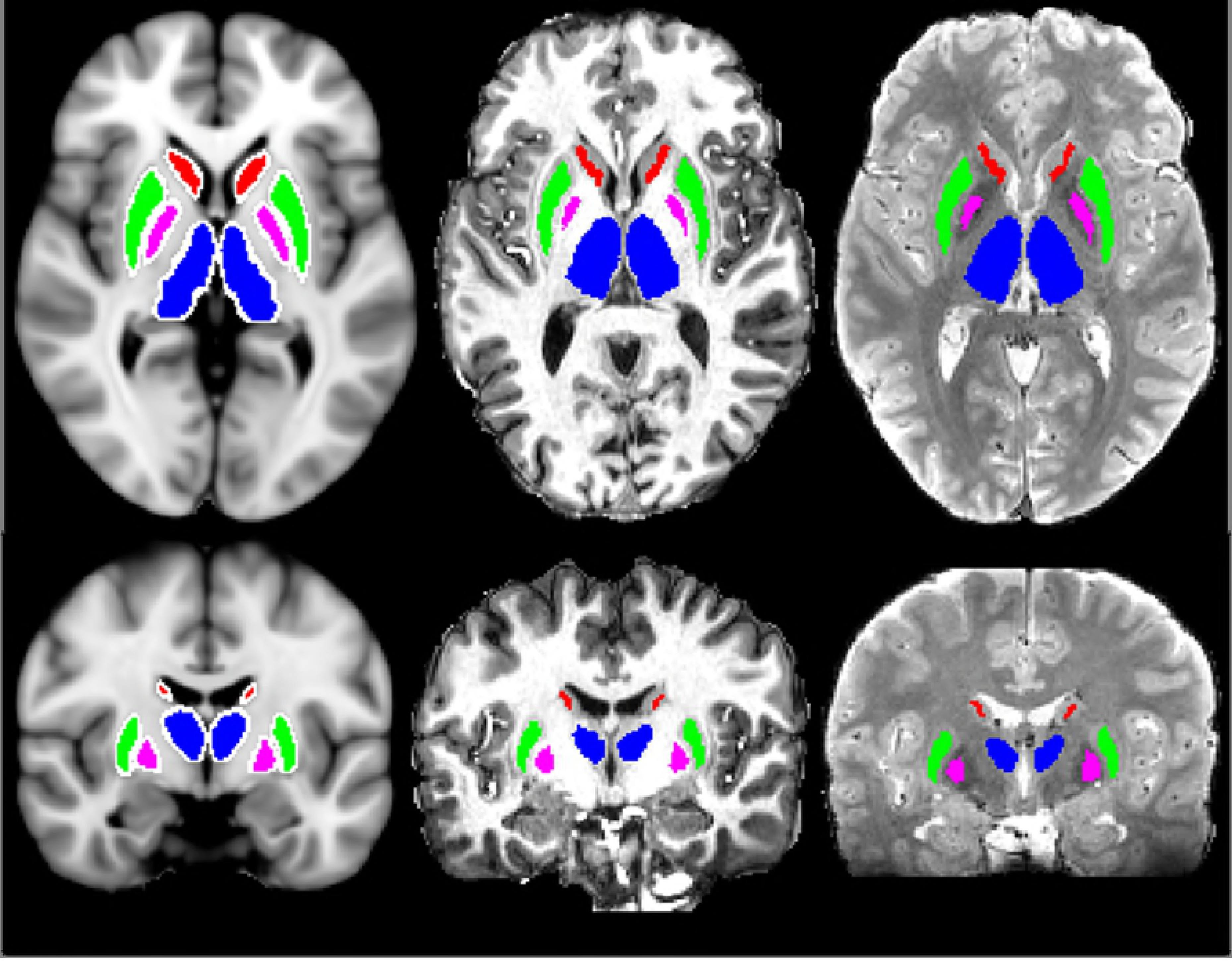
Example of structural labels in a representative participant. Axial (top row) and coronal (bottom row) views of four bilateral (thus 8) structural labels in three spaces: MNI (left), MPRAGE (center), and GRE (right) spaces. Caudate nucleus (red, CN), putamen (green, PUT), globus pallidus (purple, GP), and thalamus (blue, TH) were bilaterally segmented using the Harvard-Oxford Atlas available in FSL. Colored structures are the results of erosion of the source structures (white) using one-voxel sized 3-D kernel.

### Data analysis

All statistical analyses and data visualization were carried out using R software package (R Core team, Vienna, Austria). P-values less than 0.05 were considered as significant for all analyses. Adjustments for multiple comparisons were not performed due to the small sample size and exploratory nature of this study.

#### Pixel-based analysis

We assessed the reliability of quantitative R2* and QSM metrics processed and collected in transformed space versus those processed in native space but collected in transformed space by comparing PV distributions and PV similarity in reference to metrics from R2* and QSM processed and collected in native space.

Since all sampled data violated the assumption of normal distribution, non-parametric methods were employed to evaluate the reliability of PVs. We selected the ‘Two-sample Kolmogorov-Smirnov test (KS-test)’ to assess the distribution equality of two compared data sets. The result of KS test is the D-statistic, which quantifies a distance between empirical distribution functions of the two compared data sets. We also used the ‘Man-Whitney U-test (U-test)’ to check differences between two compared data sets.

#### Region-of-Interest (ROI) based analysis

The purpose of this analysis was to investigate ROI-based group mean comparison. This type of comparison is usually performed to detect statistical difference between two groups, e.g. patient group versus controls. In our study, we applied this analysis to examine the reliability of group mean (of all subjects) of R2* and magnetic susceptibility values in eight structures. The mean values of R2* and QSM for each structure were calculated using PVs from the pixel-based analysis. We performed a qualitative review of mean metrics using bar plots. The ‘Paired t-test’ was used to assess the reliability of the mean metrics before and after linear coregistration.

In addition to the paired t-test, we also conducted intra-class correlation (ICC) test. ICC estimates and their 95% confidence intervals were calculated based on single measurement, absolute agreement, and two-way mixed effects model using ‘irr’ package available for R. The purpose of this ICC analysis was to assess the agreement between mean quantitative metrics for each structure acquired in GRE and MPRAGE spaces across all subjects.

#### Further analysis on the impact of interpolating source GRE on resultant QSM

Noticing that coregistration included interpolation of images during alignment of two images, we investigated the impact of the interpolation of the source GRE image on the resultant QSM by interpolating GRE image to the resolution of MPRAGE image. Another QSM was then processed in GRE space by directly interpolating to the MRPAGE space resolution for comparison. The QSM results, which were processed in its native GRE and an interpolated GRE spaces, were examined by comparing images, density plots and distance measured by D-statistic from KS-test. Further, we attempted observing the error propagation along the QSM processing chain.

## Results

### Pixel-based analysis

R2* maps were robust to both coregistration approaches. Fig 2 visualizes KS-test results by comparing the empirical cumulative distribution function (ECDF) of R2* maps. KS- and U-test results comparing PVs of R2* maps showed no significant difference caused by either GRE to MPRAGE coregistration (D-statistic ranged from 0.008670 to 0.03769, U-statistic ranged from 94387 to 16655829, all p > 0.05) or direct R2* coregistration to MPRAGE space (D-statistic ranged from 0.007509 to 0.4770, all p > 0.05, U-statistic ranged from 94889 to 16684842, all p > 0.05). Complete tables for KS- and U-tests results for R2* metrics is provided in S1 and S2 Tables.

**Fig 2.**
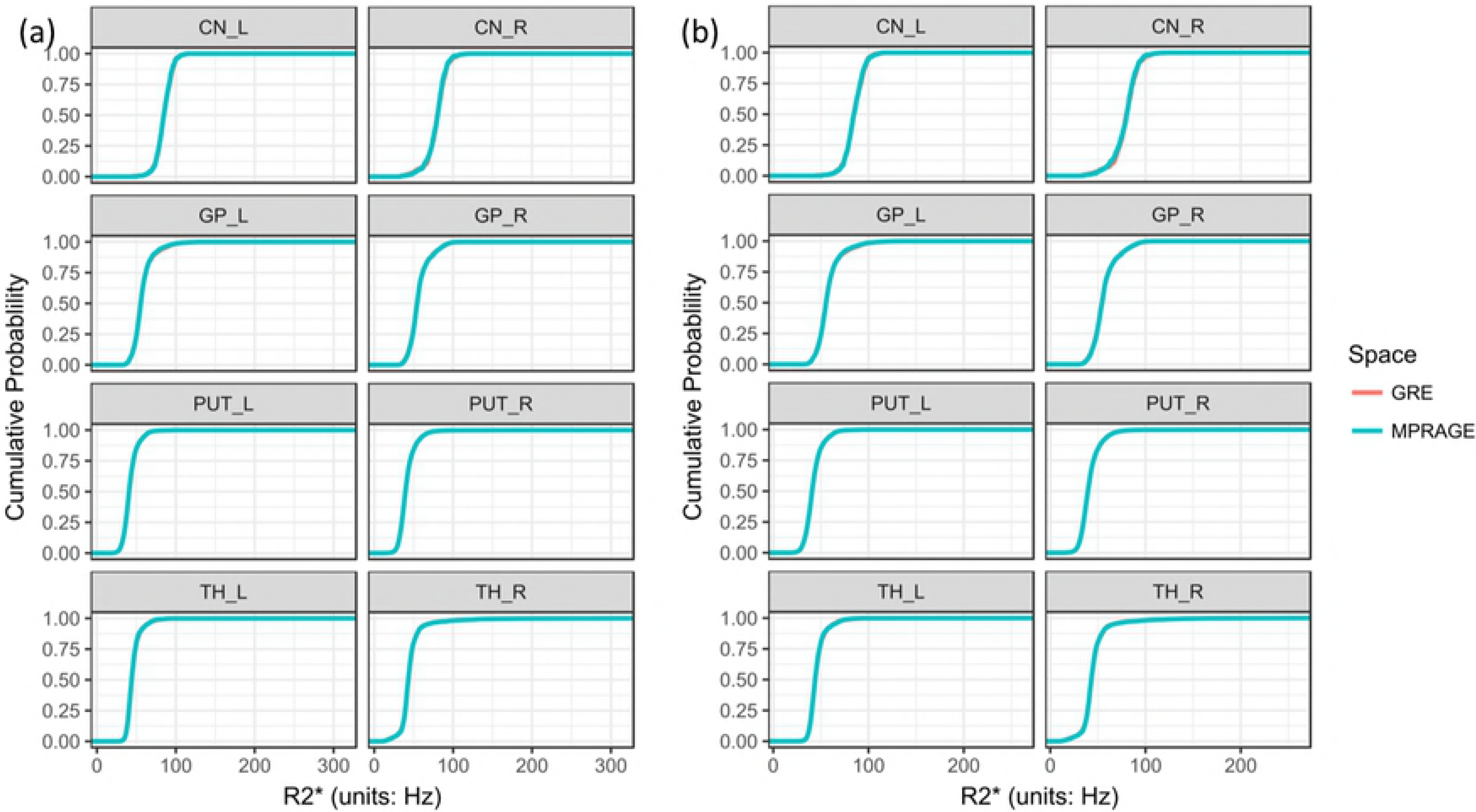
Empirical cumulative distribution functions (ECDF) of R2* maps acquired using two different coregistration approaches in a representative participant. (a) ECDF’s of R2* pixel values acquired from its native GRE space (red) and from GRE data transformed to MPRAGE space (cyan) were overlaid. In this approach, R2* calculation was performed after the coregistration of GRE data to MPRAGE space. (b) ECDFs of R2* pixel values acquired from its native GRE space (red) and from R2* map directly coregistered to MPRAGE space (cyan) were overlaid. In this approach, R2* map calculation is performed in its native space and, then, the resultant R2* map is coregistered to MPRAGE space. All ECDFs showed excellent agreement for the distributions of PVs. Similar results were confirmed in all participants. Regardless of the differences in processing steps, ECDF’s from both methods are well overlapped without significant differences in R2* values.

Fig 3 (a) visualizes discrepancies found between PV distributions of QSMs processed in GRE and in MPRAGE spaces as poorly matched ECDFs. KS-tests detected differences in 61 out of 64 structures (8 structural labels × 8 participants). U-tests found differences in 55 out of 64 structures. A full result is provided in S3 Table. On the other hand, Fig 3 (b) visualizes good agreement between QSMs processed in GRE space and directly coregistered to MPRAGE space. KS-tests detected differences in only 1 out of 64 structures. U-tests found differences in only 1 out of 64 structures. A full table of the results is provided in S4 Table. Our KS- and U-test results showed that susceptibility measurements were significantly altered in most structures of all participants after the coregistration of GRE images to MPRAGE space. In general, these results indicate that linear coregistration of the source GRE images significantly altered the distribution and PVs of the resultant QSM, while direct coregistration of QSM to a reference space did not alter the distributions and PVs.

**Fig 3.**
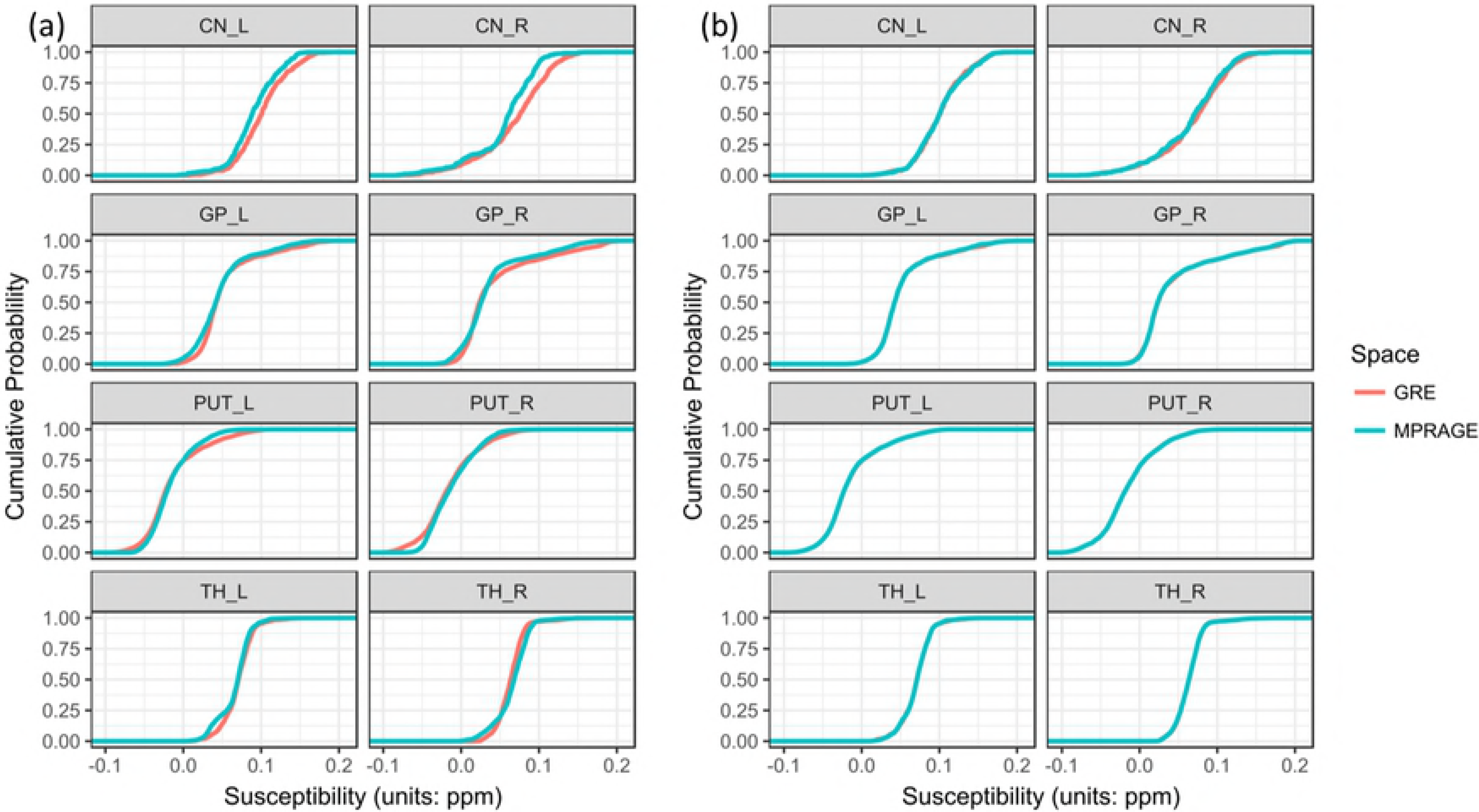
Empirical cumulative distribution functions (ECDF) of QSM by two different coregistration approaches. (a) ECDFs of magnetic susceptibility pixel values acquired from its native GRE space (red) and from GRE data transformed to MPRAGE space (cyan) were overlaid. In this approach, QSM calculation was performed after the coregistration of GRE data to MPRAGE space. (b) ECDFs of magnetic susceptibility pixel values acquired from QSM in GRE space (red) and the QSM directly coregistered to MPRAGE space (cyan) were overlaid. In this approach, QSM calculation was performed first and, then, the QSM was coregistered to MPRAGE space. Significant discrepancies were observed between QSM in GRE and MPRAGE spaces when the source GRE was transformed to MPRAGE space prior to QSM processing. On the contrary, the QSM-calculation-then-coregistration approach did not show significant differences in the pixel-based analysis except in one structure in one participant

Further analysis of frequency maps in an effort to investigate the discrepancy between QSMs acquired by two coregistration approaches was performed. Figure 4 shows one of the results in a representative participant. Noticeable differences were observed in 6 out of 8 structures between frequency maps processed in GRE and MPRAGE spaces respectively. ECDFs of frequency maps of Left and right putamen (PUT_L and PUT_R) did not show apparent discrepancies in this specific case. The differences between frequency maps did not result in a predictable matter in QSMs. Instead, corresponding QSM in PUT_L and PUT_R were significantly different between spaces where QSMs were processed as seen in Fig 3 (a). A full table of this analysis is provides in S5 Table.

**Fig 4.**
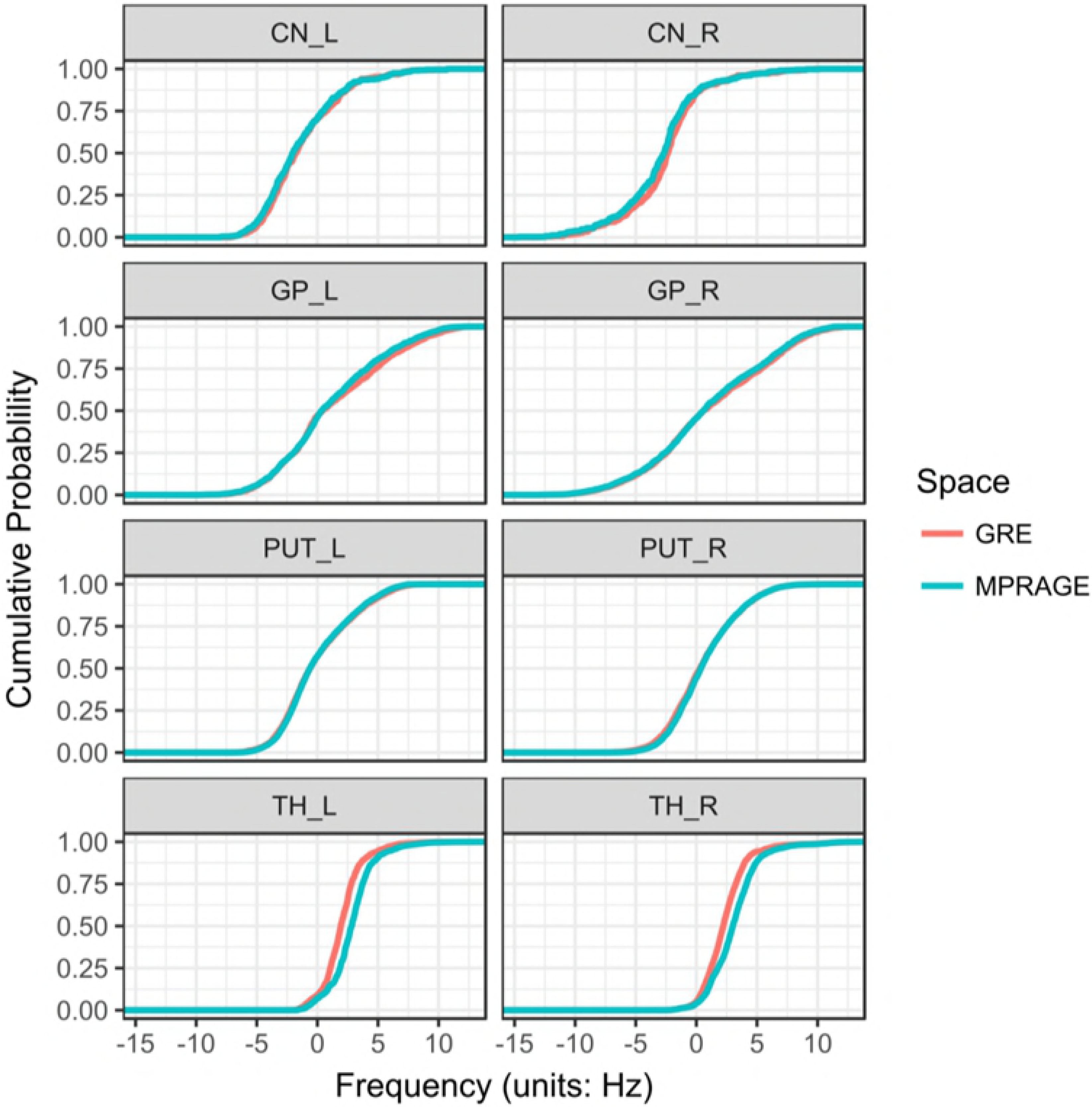
Empirical cumulative distribution functions (ECDF) of frequency maps by two different coregistration approaches. ECDFs in 8 structural labels of frequency map acquired after coregistration of source GRE data to MPRAGE space (cyan) as compared to the corresponding frequency and QSM in its native GRE space (red) in a representative participant. Relatively better overlapped ECDFs in PUT_L and PUT_R of frequency maps did not lead to better overlapped ECDFs of QSM, which still showed significant differences in the two structures as seen in Fig 3 (a).

### Region-of-Interest (ROI) based analysis

The paired t-test was conducted on the paired group-mean measurement, which is the mean (8 subjects) of means (intra-structure). The group-mean measurements of R2* were statistically equivalent (p > 0.05 for all structures) as shown in Table 1 and 2. Fig 5 shows group-mean measurements of R2* within each structure of all participants for both coregistration approaches. This indicates that group-mean R2* is reliable to either coregistration of GRE data to MPRAGE space (Fig 5 (a)) or its direct coregistration to MPRAGE space (Fig 5 (b)) in all structures.

**Fig 5.**
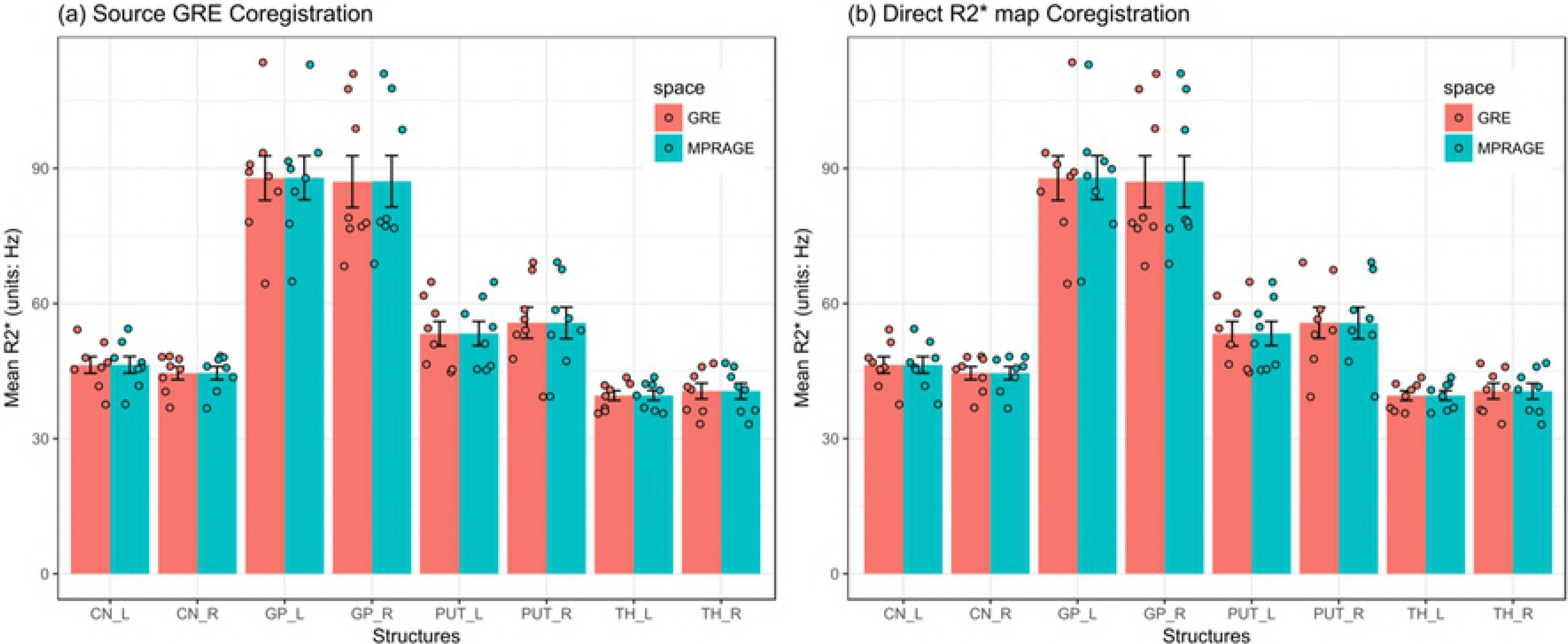
Comparison of group-mean R2* acquired by two different coregistration approaches. R2* was robust to either coregistration of source GRE data to MPRAGE space (a) or its direct coregistration to MPRAGE space (b) in all structures. See also Tables 1 and 2. (TH_L: left thalamus, TH_R: right thalamus, CN_L: left caudate nucleus, CN_R: right caudate nucleus, GP_L: left globus pallidus, GP_R: right globus pallidus, PUT_L: left putamen, PUT_R: right putamen)

**Table 1.** Mean R2* (in the unit of Hz) comparison by paired T-test for coregistration of source GRE data to MRPAGE space before R2* calculation.

| structure | mean R2* (SD)<br>in GRE space | mean R2* (SD)<br>in MPRAGE space | P-value (CI) |
| --- | --- | --- | --- |
| CN_L | 46.4 (5.22) | 46.4 (5.24) | 0.154 (-0.118, 0.0229) |
| CN_R | 44.5 (4.07) | 44.6 (4.09) | 0.776 (-0.168, 0.13) |
| GP_L | 87.8 (13.9) | 87.9 (13.7) | 0.679 (-0.494, 0.341) |
| GP_R | 87 (16.2) | 87.1 (16.1) | 0.385 (-0.284, 0.124) |
| PUT_L | 53.3 (7.68) | 53.3 (7.6) | 0.678 (-0.267, 0.185) |
| PUT_R | 55.7 (9.79) | 55.7 (9.86) | 0.747 (-0.143, 0.19) |
| TH_L | 39.5 (3.05) | 39.6 (3.04) | 0.164 (-0.0995, 0.0205) |
| TH_R | 40.6 (4.92) | 40.6 (4.96) | 0.596 (-0.0434, 0.07) |

SD: Standard deviation, CI: Confidence interval
CN\_L: left caudate nucleus, CN\_R: right caudate nucleus, GP\_L: left globus pallidus, GP\_R: right globus pallidus, PUT\_L: left putamen, PUT\_R: right putamen, TH\_L: left thalamus, TH\_R: right thalamus

**Table 2.**
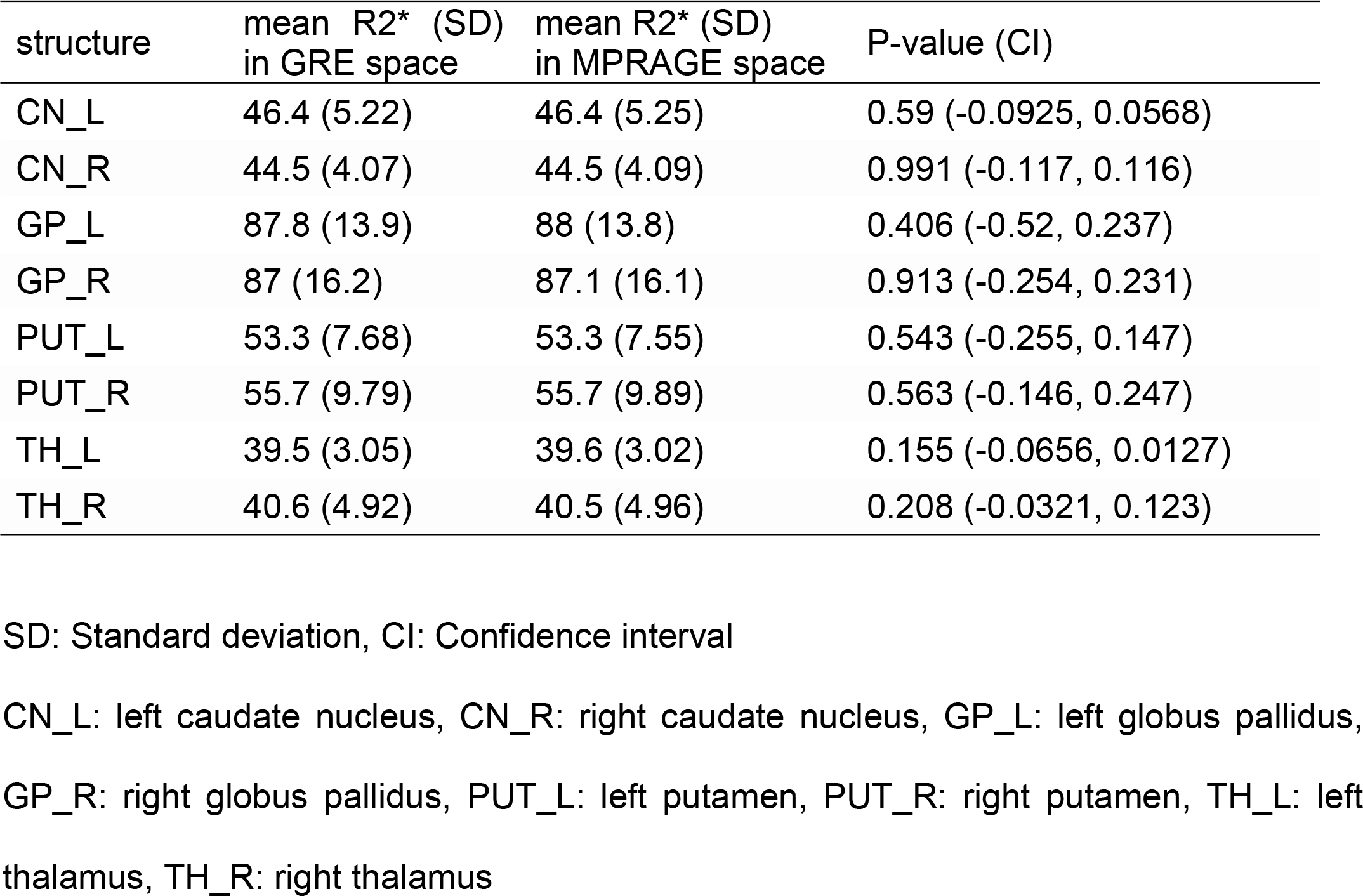
Mean R2* (in the unit of Hz) comparison by paired T-test for direct coregistration of R2* map in GRE space to MPRAGE space.

| structure | mean R2* (SD)<br>in GRE space | mean R2* (SD)<br>in MPRAGE space | P-value (CI) |
| --- | --- | --- | --- |
| CN_L | 46.4 (5.22) | 46.4 (5.25) | 0.59 (-0.0925, 0.0568) |
| CN_R | 44.5 (4.07) | 44.5 (4.09) | 0.991 (-0.117, 0.116) |
| GP_L | 87.8 (13.9) | 88 (13.8) | 0.406 (-0.52, 0.237) |
| GP_R | 87 (16.2) | 87.1 (16.1) | 0.913 (-0.254, 0.231) |
| PUT_L | 53.3 (7.68) | 53.3 (7.55) | 0.543 (-0.255, 0.147) |
| PUT_R | 55.7 (9.79) | 55.7 (9.89) | 0.563 (-0.146, 0.247) |
| TH_L | 39.5 (3.05) | 39.6 (3.02) | 0.155 (-0.0656, 0.0127) |
| TH_R | 40.6 (4.92) | 40.5 (4.96) | 0.208 (-0.0321, 0.123) |

SD: Standard deviation, CI: Confidence interval
CN\_L: left caudate nucleus, CN\_R: right caudate nucleus, GP\_L: left globus pallidus, GP\_R: right globus pallidus, PUT\_L: left putamen, PUT\_R: right putamen, TH\_L: left thalamus, TH\_R: right thalamus

The group-means of magnetic susceptibilities showed better reliability than pixel-based analyses as shown in Tables 3 and 4. Fig 6 shows comparisons of group-mean measurements of magnetic susceptibility. The results show good statistical equivalence in most structures when processed in transformed space or direct coregistration of QSMs.

**Table 3.** Mean magnetic susceptibility (in the unit of ppm) comparison by paired T-test for coregistration of source GRE data to MRPAGE space before QSM processing.

| Structure | mean $\chi_m$ (SD)<br>in GRE space | mean $\chi_m$ (SD)<br>in MPAGE space | P-value (CI) |
| --- | --- | --- | --- |
| CN_L | 0.0527 (0.02) | 0.0454 (0.0239) | 0.117 (-0.00234, 0.0168) |
| CN_R | 0.0448 (0.0155) | 0.0453 (0.0241) | 0.906 (-0.0108, 0.00975) |
| GP_L | 0.135 (0.0238) | 0.127 (0.0294) | 0.141 (-0.00362, 0.0206) |
| GP_R | 0.119 (0.039) | 0.108 (0.0447) | 0.147 (-0.00491, 0.0268) |
| PUT_L | 0.0519 (0.00817) | 0.0481 (0.0121) | 0.177 (-0.00217, 0.00976) |
| PUT_R | 0.062 (0.0167) | 0.0557 (0.017) | < 0.05 (0.00213, 0.0105) |
| TH_L | -0.00625 (0.00787) | 0.00292 (0.0147) | 0.0758 (-0.0196, 0.00124) |
| TH_R | -0.00489 (0.00992) | 0.00285 (0.0131) | 0.0775 (-0.0166, 0.00111) |

$\chi_m$ : magnetic susceptibility
SD: Standard deviation, CI: Confidence interval
CN\_L: left caudate nucleus, CN\_R: right caudate nucleus, GP\_L: left globus pallidus,
GP\_R: right globus pallidus, PUT\_L: left putamen, PUT\_R: right putamen, TH\_L: left
thalamus, TH\_R: right thalamus

**Table 4.**
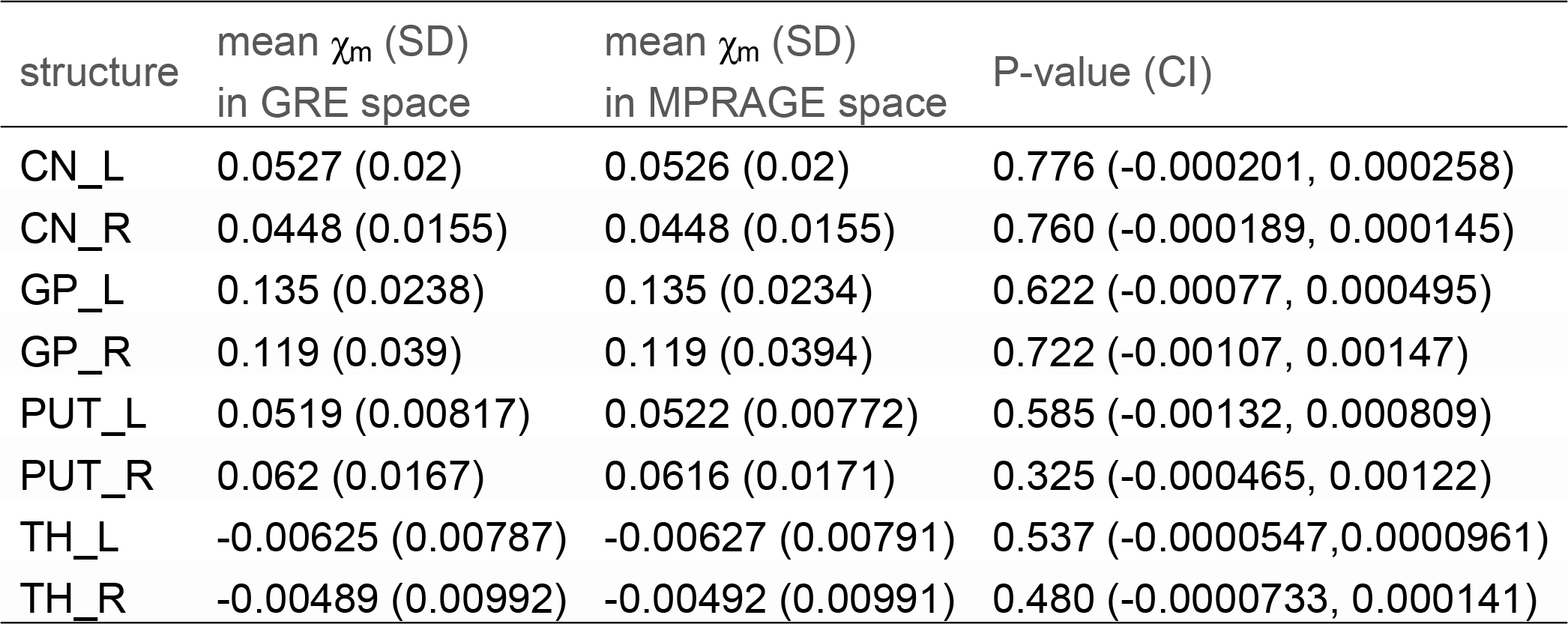
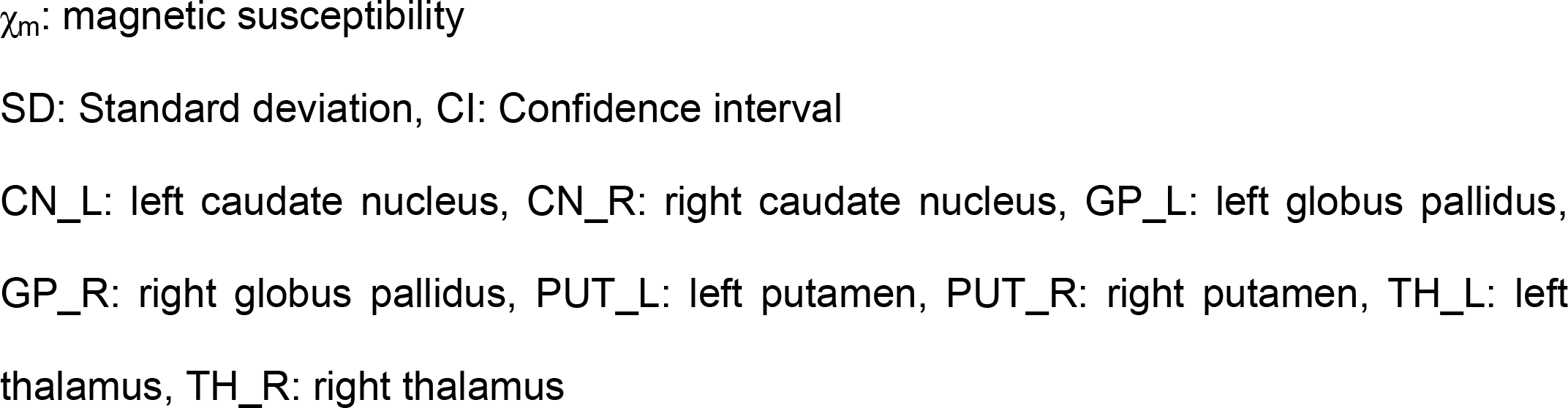
Mean magnetic susceptibility (in the unit of ppm) comparison by paired T-test for direct coregistration of QSM in GRE space to MPRAGE space.

| structure | mean $\chi_m$ (SD)<br>in GRE space | mean $\chi_m$ (SD)<br>in MPAGE space | P-value (CI) |
| --- | --- | --- | --- |
| CN_L | 0.0527 (0.02) | 0.0526 (0.02) | 0.776 (-0.000201, 0.000258) |
| CN_R | 0.0448 (0.0155) | 0.0448 (0.0155) | 0.760 (-0.000189, 0.000145) |
| GP_L | 0.135 (0.0238) | 0.135 (0.0234) | 0.622 (-0.00077, 0.000495) |
| GP_R | 0.119 (0.039) | 0.119 (0.0394) | 0.722 (-0.00107, 0.00147) |
| PUT_L | 0.0519 (0.00817) | 0.0522 (0.00772) | 0.585 (-0.00132, 0.000809) |
| PUT_R | 0.062 (0.0167) | 0.0616 (0.0171) | 0.325 (-0.000465, 0.00122) |
| TH_L | -0.00625 (0.00787) | -0.00627 (0.00791) | 0.537 (-0.0000547, 0.0000961) |
| TH_R | -0.00489 (0.00992) | -0.00492 (0.00991) | 0.480 (-0.0000733, 0.000141) |

$\chi_m$ : magnetic susceptibility
SD: Standard deviation, CI: Confidence interval

**Fig 6.**
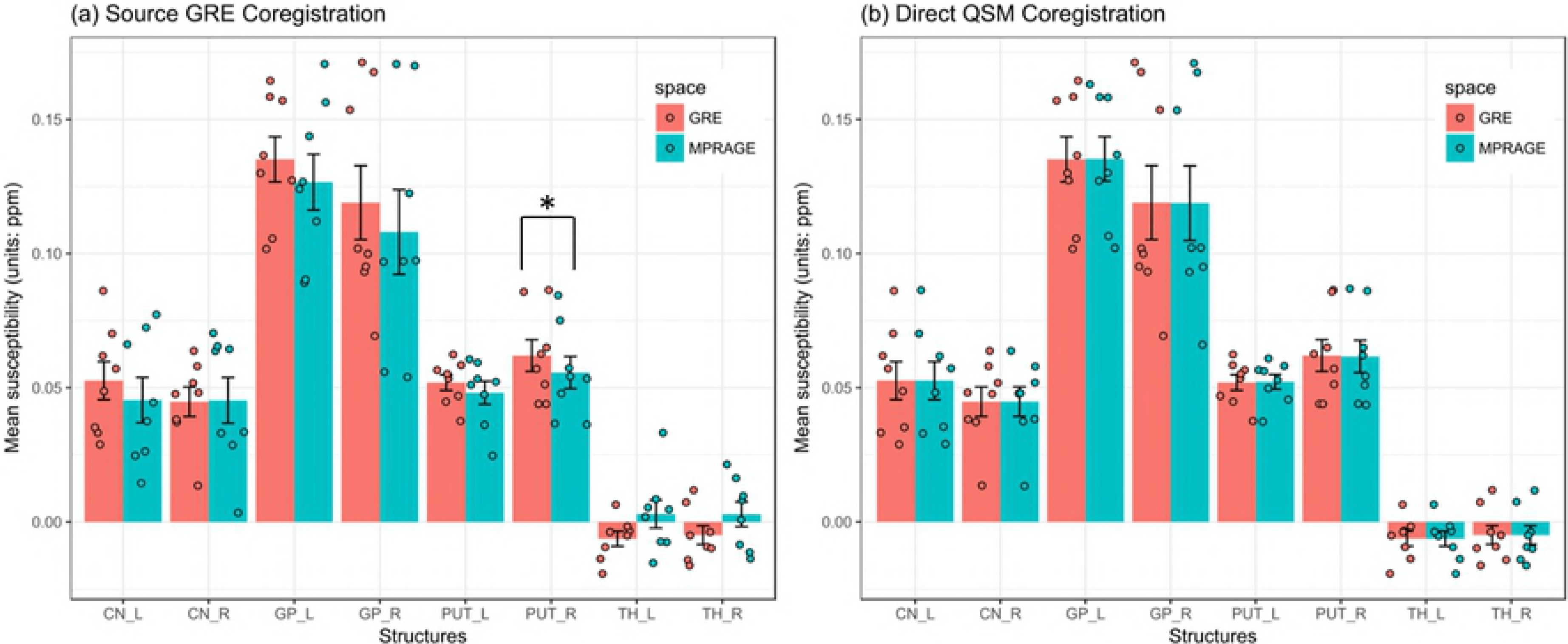
Comparison of group-mean magnetic susceptibilities acquired by two different coregistration approaches. (a)Comparison of QSM values in its native space (red) and in MPRAGE space (cyan) obtained by coregistration of source GRE data. Although differences are noticeable in group-mean susceptibility they are statistically significant except for one label (PUT_R). (b)QSM was robust to its direct coregistration to MPRAGE space in all structures. The group-mean susceptibilities were almost identical in GRE and MPRAGE spaces when QSM was directly coregistered. Therefore, coregistration of resultant QSM to a target space would be a better reproducible approach in most brain structures. (TH_L: left thalamus, TH_R: right thalamus, CN_L: left caudate nucleus, CN_R: right caudate nucleus, GP_L: left globus pallidus, GP_R: right globus pallidus, PUT_L: left putamen, PUT_R: right putamen)

The intra-class correlation coefficient (ICC) was conducted on the mean (intra-structure) measurement. For all pairs of measurements of R2*, the estimated ICCs were higher than 0.9 as shown in Table 5. For magnetic susceptibility, ICCs were higher than 0.9 between the measurements processed in GRE space and those directly transformed to MPRAGE space. ICCs were between 0.75 and 0.9 in five out of eight structures between the magnetic susceptibilities estimated in GRE space and the ones in transformed space and three structures showed ICCs between 0.5 and 0.75 or lower than 0.5.

**Table 5.** Intraclass correlation (ICC) coefficients.

| structure | R2* (GRE vs. MPRAGE) | | R2* (GRE vs direct coreg.) | | $\chi_m$ (GRE vs. MPRAGE) | | $\chi_m$ (GRE vs direct coreg.) | |
| --- | --- | --- | --- | --- | --- | --- | --- | --- |
|  | ICC (CI) | P_value | ICC (CI) | P_value | ICC (CI) | P_value | ICC (CI) | P_value |
| CN_L | 1 (0.999, 1) | <0.001 | 1 (0.999, 1) | <0.001 | 0.834 (0.38, 0.964) | <0.05 | 1 (1, 1) | <0.001 |
| CN_R | 0.999 (0.996, 1) | <0.001 | 0.999 (0.997, 1) | <0.001 | 0.834 (0.357, 0.965) | <0.05 | 1 (1, 1) | <0.001 |
| GP_L | 0.999 (0.997, 1) | <0.001 | 0.999 (0.998, 1) | <0.001 | 0.826 (0.378, 0.962) | <0.05 | 1 (0.998, 1) | <0.001 |
| GP_R | 1 (0.999, 1) | <0.001 | 1 (0.999, 1) | <0.001 | 0.879 (0.523, 0.974) | <0.001 | 0.999 (0.997, 1) | <0.001 |
| PUT_L | 0.999 (0.997, 1) | <0.001 | 1 (0.998, 1) | <0.001 | 0.734 (0.194, 0.939) | <0.05 | 0.988 (0.945, 0.998) | <0.001 |
| PUT_R | 1 (0.999, 1) | <0.001 | 1 (0.999, 1) | <0.001 | 0.897 (0.146, 0.982) | <0.05 | 0.998 (0.992, 1) | <0.001 |
| TH_L | 1 (0.998, 1) | <0.001 | 1 (0.999, 1) | <0.001 | 0.357 (-0.2, 0.807) | 0.114 | 1 (1, 1) | <0.001 |
| TH_R | 1 (1, 1) | <0.001 | 1 (0.999, 1) | <0.001 | 0.501 (-0.107, 0.867) | 0.0536 | 1 (1, 1) | <0.001 |

R2\* is in the unit of Hz
$\chi_m$ : magnetic susceptibility in ppm
CI: confidence interval
CN\_L: left caudate nucleus, CN\_R: right caudate nucleus, GP\_L: left globus pallidus, GP\_R: right globus pallidus, PUT\_L: left putamen, PUT\_R: right putamen, TH\_L: left thalamus, TH\_R: right thalamus

### Impact of coregistration of source GRE on resultant QSM

#### Impact of interpolation

Interpolation to different image resolution did not change magnitude or raw phase images visually as in Fig 7. While interpolated QSM after processing did not show significant differences (KS-test, D = 0.00175, p > 0.05) from the QSM in native space, QSM calculated from magnitude and phase images interpolated to different spatial resolution showed significant differences (D = 0.00992, p < 0.001) as in Fig 8.

**Fig 7.**
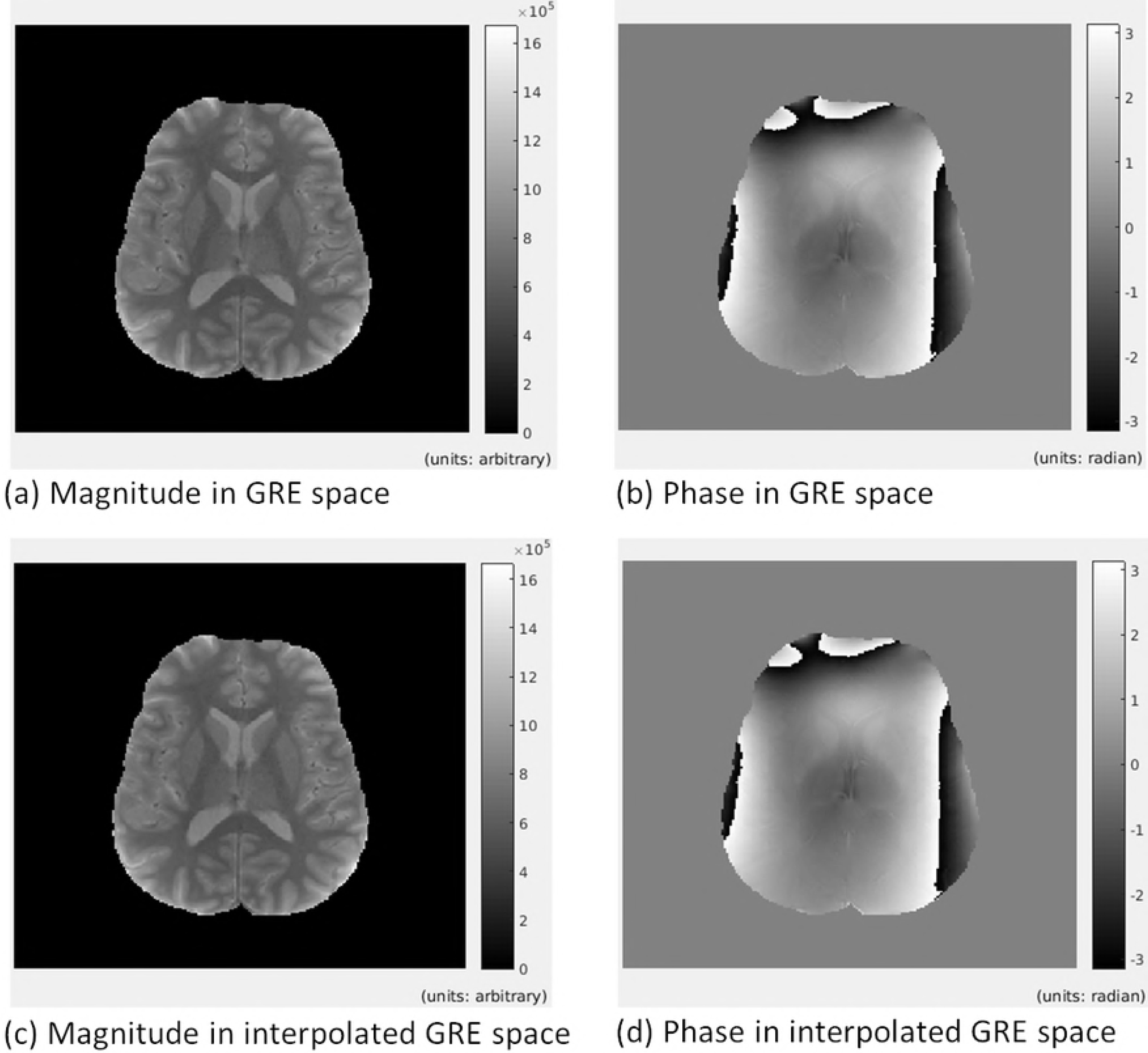
No significant impact of interpolation on raw images. Interpolation did not change the source image drastically: (a) magnitude image in GRE space (0.982×0.982 mm^2^), (b) phase image in GRE space, (c) magnitude image interpolated to MPRAGE space resolution (1×1 mm^2^ in plane resolution), (d) phase image interpolated to MPRAGE space resolution.

**Fig 8.**
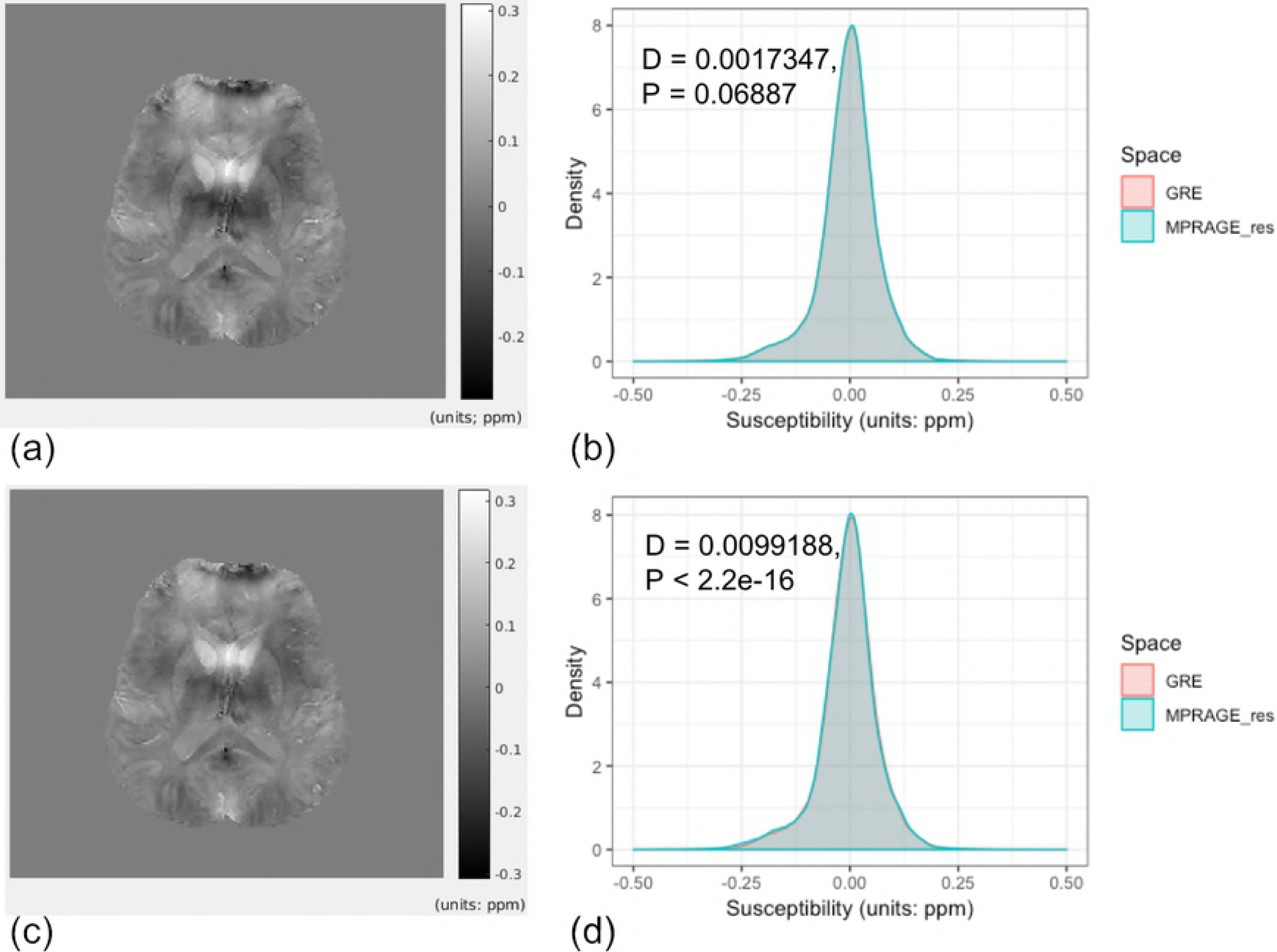
Impact of two interpolation approaches on QSM. QSM directly interpolated was not significantly different from QSM processed in its native space (a) and (b). On the contrary, source GRE interpolation resulted in significantly different QSM distributions (c) and (d).

#### Impact of coregistration on each step of QSM calculation chain

Coregistration, which included not only interpolation but also angle changes between GRE and MPRAGE space, also did not cause significant statistical difference (D = 0.000639, p = 0.977) between raw phase distributions of GRE and interpolated GRE images (Fig 9 (a)). However, when they were unwrapped, distribution was (D = 0.268, p < 0.001) significantly different (Fig 9 (b)). This difference was reflected on frequency maps (D = 0.00659, p < 0.001) (Fig 9 (c)) and eventually caused significant difference (D = 0.0212, p < 0.001) between resultant QSMs. (Fig 9 (d)) QSM from source GRE coregistration was different than source GRE interpolation from the QSM in native space.

**Fig 9.**
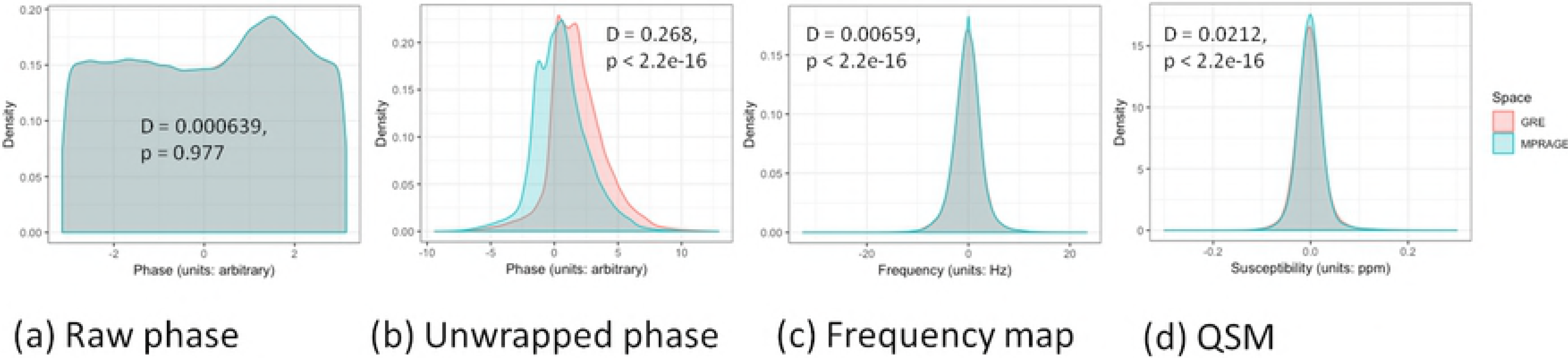
Small error in source GRE coregistration propagates along the QSM processing chain. Although coregistration did not cause statistical differences in raw phase images (a), when it was unwrapped differences were distinguishable (b). This caused differences in frequency maps and eventually caused differences in QSM (d). Phase images of 4-th echo out of 5 echoes used to calculate the frequency map and QSM are shown here.

## Discussion

### Reliability of R2* and QSM at 7T

Our primary purpose was to evaluate the impact of linear coregistration methods on GRE-derived metrics (R2* and QSM) acquired at 7T. Our results revealed that R2* was a reliable metric when processed in either native GRE space or non-native MPRAGE space, whereas magnetic susceptibility was not. The well-matched ECDFs indicate that neither linear coregistration of GRE data nor direct coregistration of R2* map to MRPAGE space alter the pixel values and distribution of R2* metrics. On the contrary, ECDFs of QSM were only well matched for direct QSM registration to MPRAGE space. This result implies that different approaches of coregistration may affect the pixel values and distribution of resultant QSM significantly in different manner. Therefore, QSM studies, in particular those employing pixel-based analysis, should be limited to susceptibility values if QSMs are calculated in native space.

### Difference between R2* and QSM processing

There are a number of possible reasons for the difference in the reliability of R2* versus magnetic susceptibility found in this study. One source of this discrepancy may arise from the intrinsically different characteristics of calculations used for R2* and QSM. R2* is calculated only from the magnitude of GRE images, while QSM (V-SHARP and SFCR in this study) utilizes mainly the phase. R2* calculated in a voxel-wise manner is an inherently local entity, while QSM reconstruction involves complicated dipole inversion that needs to take phase/frequency values in neighboring voxels into account. Since R2* maps are calculated in a voxel-wise manner from magnitude components across echo-times (TEs), any small magnitude difference within a voxel stay local without propagation to other voxels. On the contrary, phase components are determined by susceptibility sources not only within a voxel but also in surrounding and remote voxels. Elkady et al.’s work demonstrated the requirement of an extended field-of-view (5-fold) to minimize errors in QSM reconstruction of selected anatomical structure. [23] From a different perspective, their work also suggests that any alteration in QSM PV of a certain voxel could be caused by alterations in PVs of GRE phase/frequency within surrounding or remote voxels even though the PV of phase/frequency does not change in that voxel itself. This suspicion was to some extent confirmed in that alteration patterns in PVs of frequency maps did not result in similar alteration patterns in QSM PVs.

Another source of the discrepancy between our findings for R2* and QSM could be the multiple-step nature of the QSM processing used in this study (phase unwrapping, background phase removal, and dipole inversion, [24–26]), which may amplify the small differences induced by coregistration. In order to minimize such error prorogations in QSM processing, recently proposed single-step QSM methods would be preferred in future studies. [27–29]

In our study, to ensure equivalent brain masks were used in both the native GRE space and the MPRAGE spaces for background removal and QSM calculation, we used the brain mask acquired in its native space, then, applied the GRE-to-MPRAGE transformation matrix for the QSM reconstruction in the transformed space. However, if the coregistration is not perfect with small discrepancies in the brain masks over the two spaces, it could lead to differences in the estimated local frequency map during the background field removal step (as in Figure 4) and further lead to discrepancies in the final QSMs through the complicated dipole inversion step. Though the ill-posedness in QSM dipole inversion [30, 31] could be overcome by applying the calculation of susceptibility using multiple-orientation sampling (COSMOS), [32] only single orientation data was acquired for the present study, and we had to use single-orientation QSM method with regularization.

### Difference in two coregistration approaches

There is an intrinsic difference between our two coregistration approaches with previous works on reliability or reproducibility of R2* and QSM [1–4]. Previous works compared two data of an identical image resolution. However, our reliability test was performed for data sets of close but different image resolutions. From this perspective, R2* calculation was not affected when it was processed with either native resolution or non-native resolution. This indicates that a slight alteration of image resolution in a transformed space does not affect resulting PVs or their distribution in R2* maps. On the contrary, a similar reliability was not confirmed for QSM reconstruction in our study.

Previous studies have shown that direct coregistration of QSM was reliable for various transformations including intra-individual coregistration [3] and spatial normalization to MNI space. [1] Our study adds to this body of literature by providing evidence of discrepancies introduced by processing QSM in a non-native space. Collectively, these results suggest that QSM must be processed before coregistration of its source GRE image to another image space. On the other hand, our results showed statistical invariance of R2* metrics after the coregistration of GRE images at both pixel and group levels. In these reliability tests, the agreement of R2* measurements before and after coregistration suggest that R2* maps may serve as robust measurements regardless of the timing of coregistration.

In contrast to the robustness of R2* metric to different coregistration approaches at both pixel- and ROI-levels, differences observed in pixel-based analyses of QSM measures were not statistically significant in ROI-based analyses when QSM was processed in the transformed space. This is presumably due to the fact that individual differences at the pixel level were averaged out for the ROIs. Although group mean values of susceptibilities of most structures were statistically identical, noticeably bigger differences were observed when QSM was processed in the transformed space as compared to directly coregistered QSM values consistent with the pixel-value based analysis. This result also suggests that ROI-based analysis and pixel-based analysis of QSM may lead to contradictory conclusions – especially when QSM was derived from transformed GRE data instead of its native space.

In addition, paired-t-test (for the paired χ_m_ mean of means) and ICC (for grouped intra-structure means of χ_m_) showed unreliability in different structures. This indicates that a chosen statistical analysis approach may also affect the interpretation of a study for group comparison when QSM is processed in a non-native space.

### Impact of interpolation and coregistration

As an essential process in any coregistration, interpolation appeared to be a key factor affecting the QSM result in a non-native space. It is closely related to what source phase values are fed into the dipole inversion processing chain. Because of the ill-posed nature of dipole inversion, slightly different source values as well as voxel counts caused by altered voxel dimensions may lead to evidently different QSM after a long chain calculation pipeline. Our results clearly showed how slightly different voxel dimension of a source image could lead to distinguishable differences in resultant QSM. Furthermore, these results suggest when large-scale cross-site data sets are compared, it is critical to keep the image parameters as close as possible with extra care on source image resolution in voxel based analyses.

### Limitations

There are a few limitations in our study. Although our results showed excellent statistical invariance of R2* to the linear coregistration of its source GRE image at the pixel and group levels, the reliability of R2* for a non-linear transformation of the source GRE images was not tested. Hence, our finding has to be limited to linear coregistration. The robustness of statistical inference derived from R2* maps and QSMs was not verified yet, and it is beyond the scope of this study. Our future work will include the reliability of statistical relationships between quantitative metrics and clinical measures under a similar coregistration of GRE images in a larger cohort including control and patient groups. Further, our data were single-orientation QSM’s. Therefore, we could not verify if our results would apply for multi-orientation QSM, such as COSMOS.[32] However, it has to be noticed that multi-orientation QSM increases total scan time and, hence, is often impractical in a clinical study. [25] Furthermore, in our study, QSM calibration to a reference region was not performed. Although QSM calibration is being employed in recent QSM studies, there is no agreed reference region for this referencing purpose.[1, 33–36] CSF has frequently been used for the reference region in many studies.[33, 34] However, other regions have also been used. [35] Some studies did not perform QSM calibration at all. [1, 36] There have also been efforts to find suitable reference tissues for QSM. [35, 37] However, the selection of reference region can be problematic when the region includes pathologic tissues. Because we focused on the assessment of R2* maps and QSM before and after the source GRE image coregistration in the context of reliability or reproducibility instead of the accuracy of the metrics, analysis without QSM calibration to a reference region should not undermine our study results. We did not conduct further analyses on unwrapping, background removal and QSM reconstruction algorithms because it is beyond the scope of our study.

## Conclusions

To the best of our knowledge, the current study is the first report providing quantitative analysis of the influence of source GRE image transformation on the resultant parametric maps, R2* map and QSM in particular. Our study is in line with and support prior reliability studies, but is distinguished from other work in that coregistration was performed for not only resultant parametric maps but also source GRE images providing complementary data which have not been explored before. In conclusion, R2* maps are reliable when processed from GRE data linearly transformed to a non-native space, whereas QSM is not. Hence, caution is advised when using QSM in a multi-modal MRI study, and it is strongly recommended to process QSM in its native space prior to any coregistrations.

## Acknowledgment

The support of many team members makes work like this possible. We would like to acknowledge the support of the staff of the F.M. Kirby Center, including MRI technologists Terri Brawner, Kathleen Kahl, and Ivana Kusevic and the Center’s director, Dr. Peter van Zijl. We would also like to acknowledge the support of study coordinators Ms. Julie Fiol and Kerry Naunton.

